# Convergence Assessment for Bayesian Phylogenetic Analysis using MCMC simulation

**DOI:** 10.1101/2021.05.04.442586

**Authors:** Luiza Guimarães Fabreti, Sebastian Höhna

## Abstract

1. Posterior distributions are commonly approximated by samples produced from a Markov chain Monte Carlo (MCMC) simulation. Every MCMC simulation has to be checked for convergence, i.e., that sufficiently many samples have been obtained and that these samples indeed represent the true posterior distribution.
2. Here we develop and test different approaches for convergence assessment in phylogenetics. We analytically derive a threshold for a minimum effective sample size (ESS) of 625. We observe that only the initial sequence estimator provides robust ESS estimates for common types of MCMC simulations (autocorrelated samples, adaptive MCMC, Metropolis-Coupled MCMC). We show that standard ESS computation can be applied to phylogenetic trees if the tree samples are converted into traces of absence/presence of splits.
3. Convergence in distribution between replicated MCMC runs can be assessed with the Kolmogorov-Smirnov test. The commonly used potential scale reduction factor (PSRF) is biased when applied to skewed posterior distribution. Additionally, we analytically derive the expected difference between split frequencies (EDSF) and show that it depends on the true frequency of a split. Hence, the average standard deviation of split frequencies is too simplistic and the EDSF should be used instead to check for convergence in split frequencies.
4. We implemented the methods described here in the open-source R package Convenience (https://github.com/lfabreti/convenience), which allows users to easily test for convergence using output from standard phylogenetic inference software.

## Introduction

In the last two decades, Bayesian inference has become a widely used framework for performing statistical analyses in phylogenetics and macroevolutionary biology (Nascimento *et al.*, 2017; Lartillot, 2020). The goal of a Bayesian analysis is to compute the posterior probability distribution of the parameters given the observed data. Unfortunately, we cannot compute this posterior distribution analytically for virtually all realistic and empirically interesting models. Instead, one often applies sampling based methods such as Markov chain Monte Carlo sampling (MCMC, Metropolis *et al.*, 1953; Hastings, 1970). MCMC algorithms produce (autocorrelated) samples from the desired posterior distribution. Thus, the frequency of parameter samples corresponds to their posterior probability. As with any stochastic sampling based method, the researcher needs to make sure that (a) sufficiently many samples have been obtained, and (b) the samples are indeed representative of the posterior distribution. The process of determining whether these conditions have been met is called *convergence assessment*.

Convergence assessment is a widespread problem for Bayesian analyses using MCMC methods (although one should check for convergence when using any stochastic search algorithms). Every class and lecture about MCMC methods teaches practitioners to check for convergence but in practice these checks are neither standardized nor consistently performed (Harrington *et al.*, 2021). Unfortunately, traditional MCMC convergence assessment methods from the statistical community have several shortcomings when applied to Bayesian phylogenetics. First, phylogenetic trees are a very peculiar and difficult type of parameter for which common convergence tests that assume continuous parameter values cannot be applied. Thus, specific methods which transform phylogenetic trees into distances have been proposed (e.g., Lanfear *et al.*, 2016). Second, all widely used convergence assessment approaches used in phylogenetics require manual interaction, often through visual inspection (Nylander *et al.*, 2008; Warren *et al.*, 2017; Rambaut *et al.*, 2018). Visual inspection renders convergence assessment irreproducible by reviewers and other researchers. Moreover, visual inspection makes it unfeasible to apply convergence assessment for genomic datasets with thousands of parameters (e.g., each gene tree and gene-specific substitution model parameters) and for simulation studies with hundreds or thousands of MCMC runs.

In this manuscript we aim to develop a convergence assessment approach for phylogenetics that fulfills the following criteria: (i) it checks whether a single MCMC run needs to be run longer; (ii) it compares multiple independent MCMC runs to check if one of the runs got trapped in an area of parameter space and thus did not sample from the target posterior distribution; (iii) it uses statistically motivated and mathematically derived thresholds, (iv) with longer MCMC runs the chance of convergence increases towards one and does not plateau at a 5% rejection level; and (v) it can be applied without manual interactions. Our motivation and final goal is to provide a tool that, if used and its output provided in publications that used MCMC simulations in phylogenetics, we as readers or reviewers could easily verify if convergence was achieved.

## Methods

As we stated above, convergence assessment consist of checking whether (1) enough samples have been obtained, and (2) if these samples represent the true posterior distribution. The first aspect concerns the *precision* of the estimators (e.g., the posterior mean or the 95% credible interval). A common question is: “Has the MCMC simulation run long enough?” Or in other words, “Would more samples and/or a longer MCMC run change the estimates?” The most frequent approach to answer this question in phylogenetics is to assess whether the effective sample size (ESS) is larger than 200 (Rambaut *et al.*, 2018). That is, if samples obtained from the chain are equivalent to 200 or more independent samples from the posterior distribution, then one assumes that the MCMC has been run long enough. However, this threshold of 200 samples is arbitrary and without any statistical motivation, as stated in Rambaut *et al.* (2018). Here, we turn the question around and ask instead “How many samples do I need to obtain a sufficiently precise estimate?” We will derive the number of effective samples needed to obtain a specified precision. We assume, following standard statistical practice, that our parameter estimates will not change once this precision has been reached (Gong & Flegal, 2016). Thus, as suggested by Rambaut *et al.* (2018), a single MCMC run has been run sufficiently long if the ESS is larger than our derived threshold value. So far, the ESS for convergence assessment is primarily applied to continuous model parameter but not phylogenetic trees (but see Lanfear *et al.*, 2016; Warren *et al.*, 2017). Thus, we will develop and test a new method to compute the ESS for splits of a phylogeny.

The second aspect concerns the *reproducibility* of the stochastic sampling algorithm. It could be the case that the MCMC simulation got stuck in some area of the parameter space and never converged to the true posterior distribution. Therefore, we will adopt a test that compares samples from two independent MCMC simulations. In phylogenetics, the often used approach to compare samples from two independent MCMC simulations is the potential scale reduction factor (PSRF, Gelman & Rubin, 1992) and the average standard deviation of split frequencies (ASDSF, Lakner *et al.*, 2008). Neither the PSRF nor ASDSF have clear and statistically motivated thresholds. Furthermore, the PSRF is very dependent on the shape of the posterior distribution (see Supplementary Information section S2). Therefore, the PSRF does not fulfill our criteria of a robust convergence assessment method that would eventually accept the samples if the MCMC were run sufficiently long and sampled from the true distribution. Instead, we propose to use the Kolmogorov-Smirnov test (KS-test, Kolmogorov, 1933; Smirnov, 1939). The ASDSF also has shortcomings which we will address here.

### PRECISION OF AN ESTIMATOR TO ASSESS SUFFICIENTLY MANY SAMPLES

Ideally we would like to compute our parameter estimates as precisely as possible. However, in many situations, such as when the parameter estimate is computed using numerical methods, we can not obtain the estimated value with arbitrary precision. Instead, we content ourselves if the computed parameter estimates is precise to, for example, a certain number of significant digits. The number of significant digits depends on the sample variance. For example, if we want to estimate the average body size of population then we might want to have at least a precision on the scale of centimeters, but if we want instead to estimate the average flight distance of migrating birds, then a precision up to a few meters could be completely sufficient. Similarly, no one would trust the mean estimate from only a handful of observations but most people would trust an estimate of a population mean if hundreds of observation have been taken. Given these considerations, we specify the number of samples needed from an MCMC based on the desired precision.

Our threshold value for the ESS is derived from the standard error of the mean 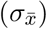, which is defined as 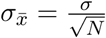 where *σ* is the standard deviation of the sample and *N* the sample size. 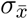 measures the error associated with the estimated mean value; or, in other words, the precision of the mean estimator. Both the variance and the sample impact how precise the mean estimate can be. For example, a sample *x* from a distribution with a larger variation has a higher error associated with the mean (or lower precision) than a sample from a distribution with lower variation. Therefore, we define an acceptable standard error to have a width smaller than or equal to 1% of the width of the 95% probability interval of the true distribution. If we assume a normal distribution as our reference distribution, then the width of the 95% probability interval is approximately equal to 4*σ* (see Figure S1). Thus, we derive a threshold value for the ESS based on the specified precision of 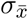 as

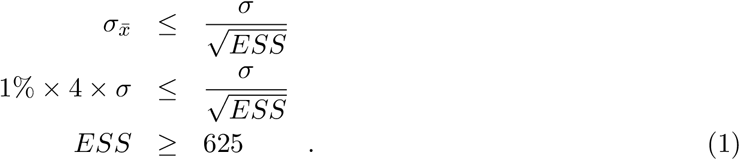

We will use this threshold value of a minimum required ESS of 625 as our reference, but other researchers could derive their own justified threshold for the ESS by specifying a different allowed standard error 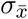 Table 1 shows some examples of the width of 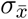 regarding the 95% probability interval and ESS values.

**Table 1:** List of minimum ESS thresholds based on precision of 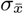.

| Width of $\sigma_{\bar{x}}$ | ESS |
| --- | --- |
| 0.5% | 2500 |
| 1% | 625 |
| 1.77% | 200 |
| 2% | 156.25 |
| 5% | 25 |

#### Assessing ESS estimation of continuous parameters

The ESS plays a fundamental role in assessing convergence. Therefore it is crucial that we can estimate the ESS correctly. The ESS can be defined as the number of samples *N* taken from an MCMC simulation divided by the autocorrelation time (*τ*)

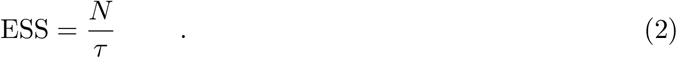

Most methods estimate the autocorrelation time *τ* instead of computing the ESS directly. Nevertheless, Equation 2 shows that both the ESS and *τ* are directly correlated. The ESS (or *τ*) can be estimated using different approaches (for an overview, see Thompson, 2010). We investigated three different commonly used ESS estimation implementations: (1) the `R` package `CODA` (Plummer *et al.*, 2006); (2) the `R` package `MCMCSE` (Flegal *et al.*, 2020); and (3) our own re-implementation in `R` of the ESS computation algorithm implemented on the software `Tracer` (Rambaut *et al.*, 2018). `CODA` estimates the autocorrelation as an auto-regressive process and estimates the spectral density at frequency 0 (Hamilton, 1994; Thompson, 2010). `MCMCSE` uses a batch means approach to estimate the autocorrelation time (Geyer, 1992; Flegal & Jones, 2010). `Tracer` uses the initial sequence estimator (Straatsma *et al.*, 1986). We will address these different methods by the names of the tools in which they were implemented (`CODA`, `MCMCSE` and `Tracer`). We test below common variants of MCMC algorithms used in phylogenetics (e.g., Metropolis-Coupled MCMC (Geyer, 1991; Altekar *et al.*, 2004) and adaptive MCMC (Haario *et al.*, 1999, 2001) as well as specific parameters such as the phylogeny).

##### Autocorrelated Samples

Samples obtained from an MCMC simulation are rarely (not to say never) drawn independently. For that reason, it is important to evaluate the ESS estimation in the case of autocorrelated samples. Unfortunately, we never know the true autocorrelation *τ* for a specific MCMC algorithm because it both depends on the implementation and data. Therefore, we developed two new algorithms that mimic an MCMC simulation and produce an autocorrelated sample with specified and known autocorrelation time *τ*. The first algorithm (Algorithm 1) generates *N iid* values and resamples these values while sampling each value at least once and keeping the order of the original values. The second algorithm (Algorithm 2) generates a sequence of *N* * *τ* samples where the (*i* + 1)th sample is either drawn from the chosen distribution with probability 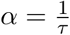 or set to the same value as the *i*-th sample with probability 1 − *α*. Although these two algorithms work differently, they produce samples with the same characteristics of autocorrelated samples by, on average, *τ* identical values consecutively. Such a behavior can be observed in MCMC samples specifically for phylogenetic trees, but also for continuous parameters when the acceptance rate is low. We included both algorithms for completeness although we present here only the results based on Algorithm 1.

**Algorithm 1.**
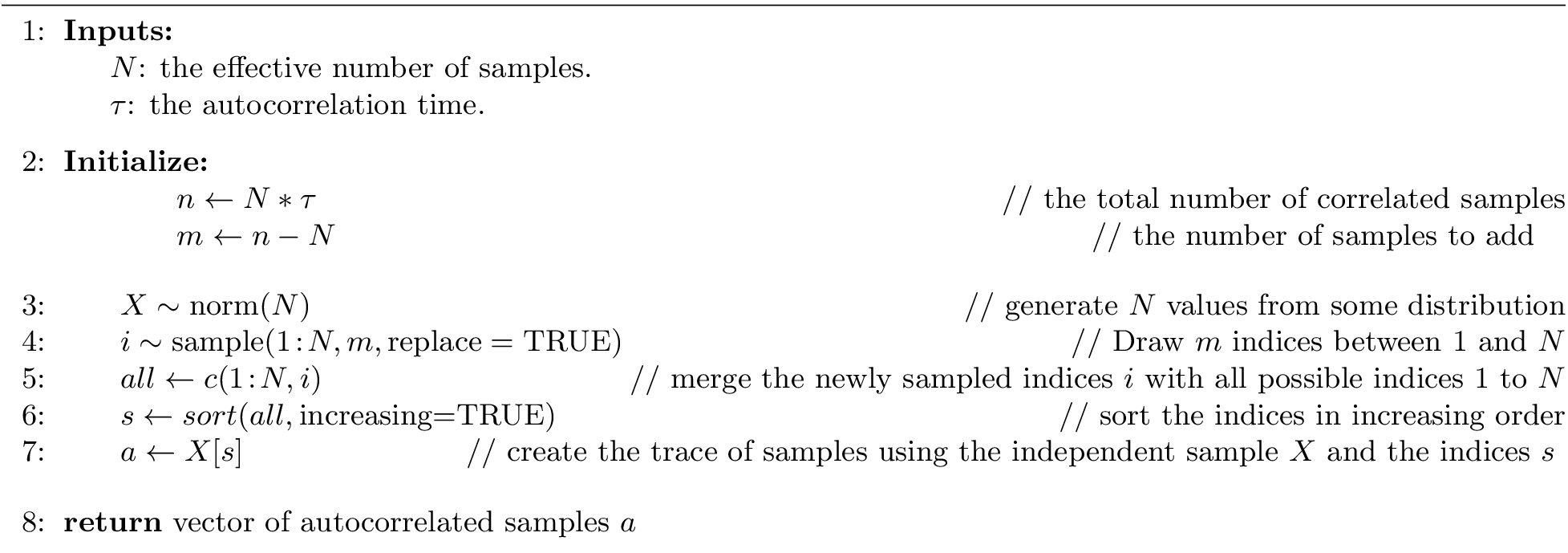
Simulating samples from an MCMC algorithm with known autocorrelation time by resampling *iid* values.

**Algorithm 2.**
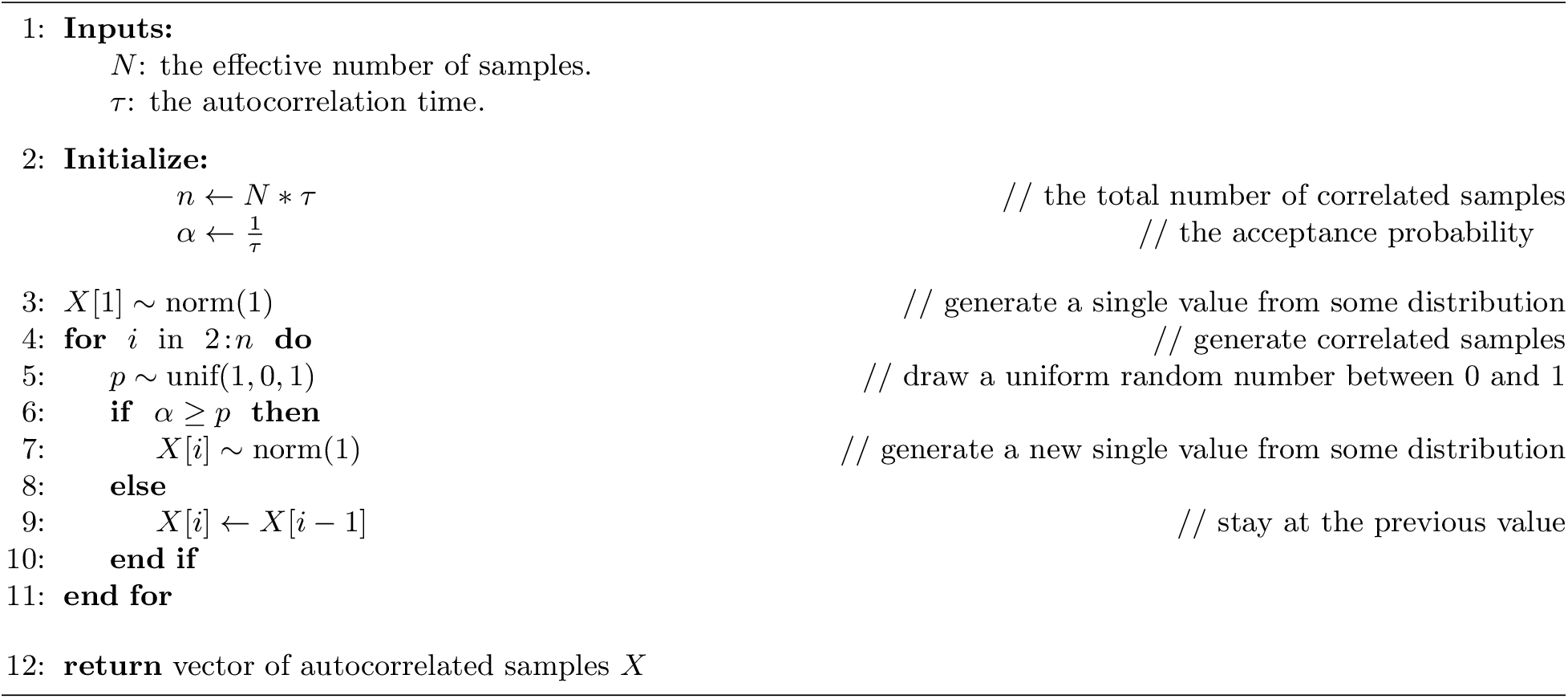
Simulating samples from an MCMC algorithm with known autocorrelation time by accepting new values with probability 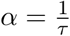.

We simulated 1,000 replicates with an ESS of *N* = {100, 200, 300, 400, 500, 625, 800, 1000} with samples drawn from a normal distribution with mean *μ* = 0 and variance *σ*^2^ = 1 and varied the autocorrelation time (ACT) between *τ* = {1, 5, 10, 20, 50, 100, 250, 500, 750, 1000}. An ACT of *τ* = 1 is equivalent to independent sampling and thus represents a baseline comparison (see Supplementary Material Section S2). Only `Tracer` produced robust ESS estimates for all values of *τ* (Figure 1). `MCMCSE` was robust to most values of the ACT, except for *τ* = 50 where the ESS were overestimated (Figure 1). Interestingly, `CODA` performed particularly bad for large *τ*. Thus, we recommend the use of `Tracer` to evaluate ESS values of MCMC samples over `MCMCSE` and `CODA`.

**Figure 1:**
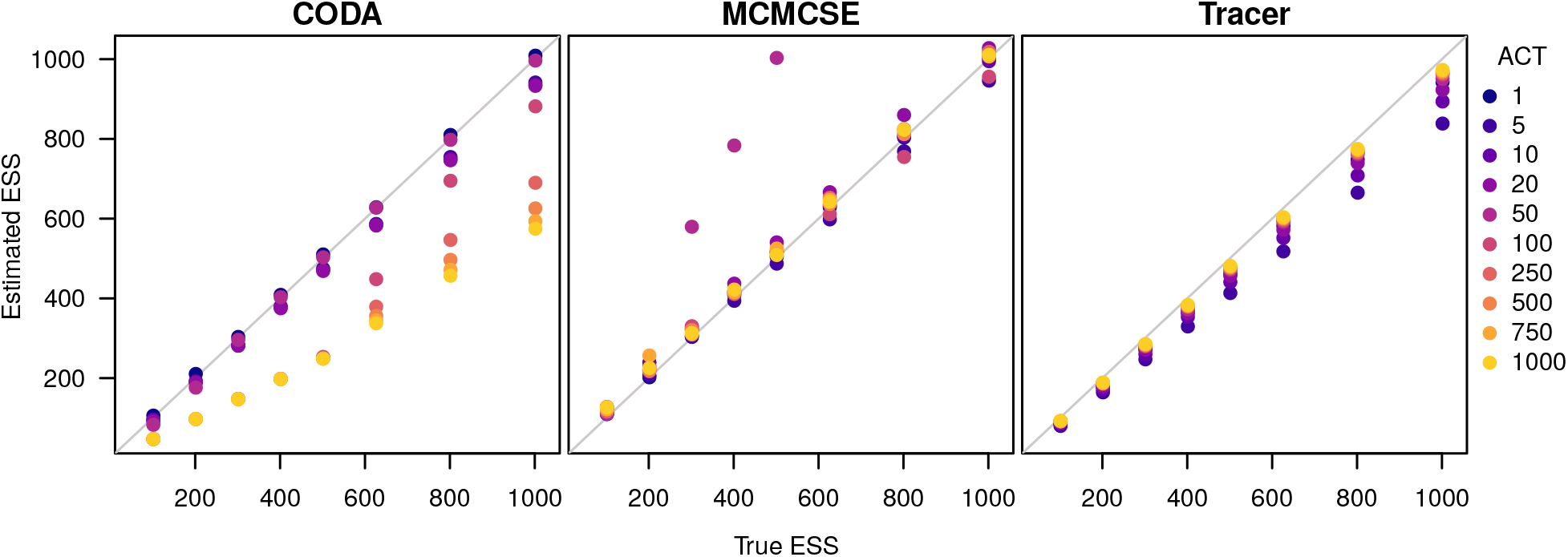
The estimated ESS for autocorrelated samples. The x-axis is the true ESS used to generate the sample, the y-axis is the estimated ESS. The left panel displays the ESS estimated using `CODA`, the central panel displays the ESS estimated using `MCMCSE` and the right panel displays the ESS estimated according to `Tracer`. The different colored dots represent different autocorrelation times. The blues dots are the cases that the samples have autocorrelation of 1. As the colors get lighter, the autocorrelation time increases.

##### Samples from Metropolis-Coupled MCMC

In Bayesian phylogenetics, Metropolis-coupled MCMC (MC^3^ or MCMCMC) is applied frequently to improve mixing of MCMC chains and thus efficiency (Altekar *et al.*, 2004; Müller & Bouckaert, 2020). An MC^3^ algorithm consists of *K* independent MCMC chains but samples are only taken from the currently active/cold chain. The MC^3^ algorithm proposes swaps between chains every *S* iterations. For more details about different MC^3^ algorithms, e.g., swap frequencies and acceptance frequencies, we refer the reader to Altekar *et al.* (2004) and Müller & Bouckaert (2020).

The samples from an MC^3^ algorithm might show different characteristics, for example, when the autocorrelation structure is seemingly broken due to swaps between chains. This behavior could mislead the estimation of the ESS. Therefore, we evaluated the ESS estimation accuracy for MC^3^ samples with different swap frequencies. First, we developed a new algorithm with a know autocorrelation *τ* to mimic MC^3^ (Algorithm 3). Our MC^3^ algorithm generates *K* independent MCMC chains using either Algorithm 1 or Algorithm 2. Then, swapping between these independent chains is performed by randomly selecting one of the chains after every *S* iterations. Note that our Algorithm 3 treats all chains as cold chains and thus always accepts a jump. However, the important feature that we want to test here is the discontinuity introduced by the jumps between chains, and if this discontinuity introduces problems for ESS estimation methods. Thus, we choose to neglect that heated chains traverse the parameter space faster than cold chains. The effect would most likely be irrelevant since our Algorithm 1 and Algorithm 2 produce independent samples once a new sample is accepted.

**Algorithm 3.**
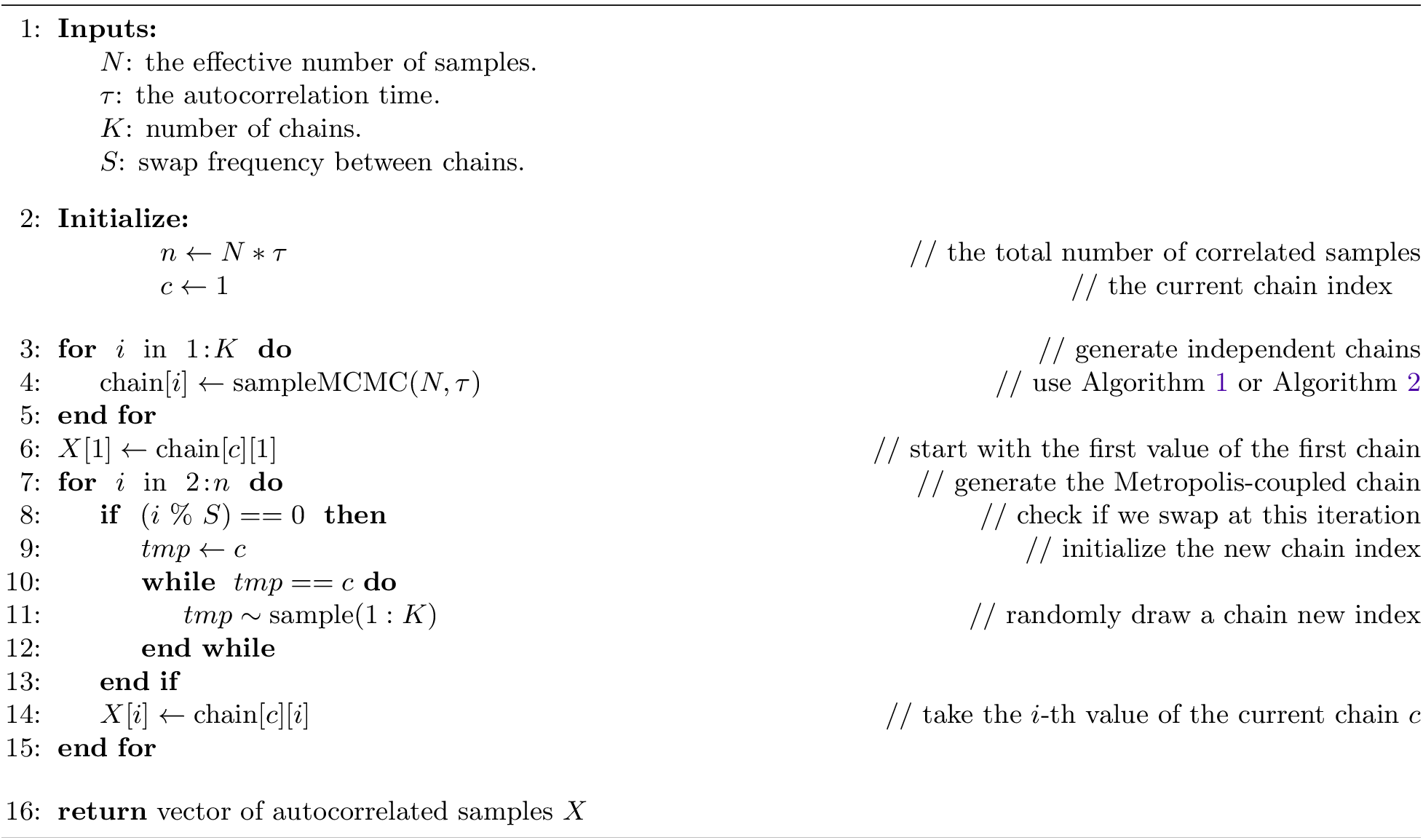
Simulating samples from an MC^3^ algorithm with known effective sample size.

In this evaluation of the three ESS estimation methods, we simulated 1,000 replicates of MC^3^ samples using Algorithm 3 with the standard number of chains *K* = 4 and an arbitrarily chosen ACT of *τ* = 20. We varied the swap frequency *S* = {1, 2, 5, 10, 20, 50, 100} in our simulations.

To derive an expectation of how many effective samples we should get using our MC^3^ algorithm, let us consider a window of *τ* samples. In our independent MCMC chains it takes, on average, *τ* iterations until we obtain a new value. We swap exactly every *S* iterations and therefore we swap 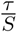times within a window of size *τ*. For every time that we swap to a chain that we did not visit yet within the window *τ* we obtain a new value. Hence, in our sample from the MC^3^ algorithm we expect that we have, on average, at least one independent value and at most *K* independent values within *τ* iterations. We derive our expectation of the ESS under the MC^3^
algorithm as

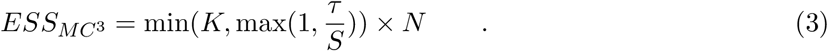

We observed that all three methods perform virtually identical for samples from our MC^3^ algorithm (Figure 2). Surprisingly, we never estimated the ESS of the MC^3^ algorithm as high as *k* × *N* (the number of chains times the ESS per chain). This, in fact, is very reassuring because it indicates that too frequent swapping will not artificially inflate ESS values.

**Figure 2:**
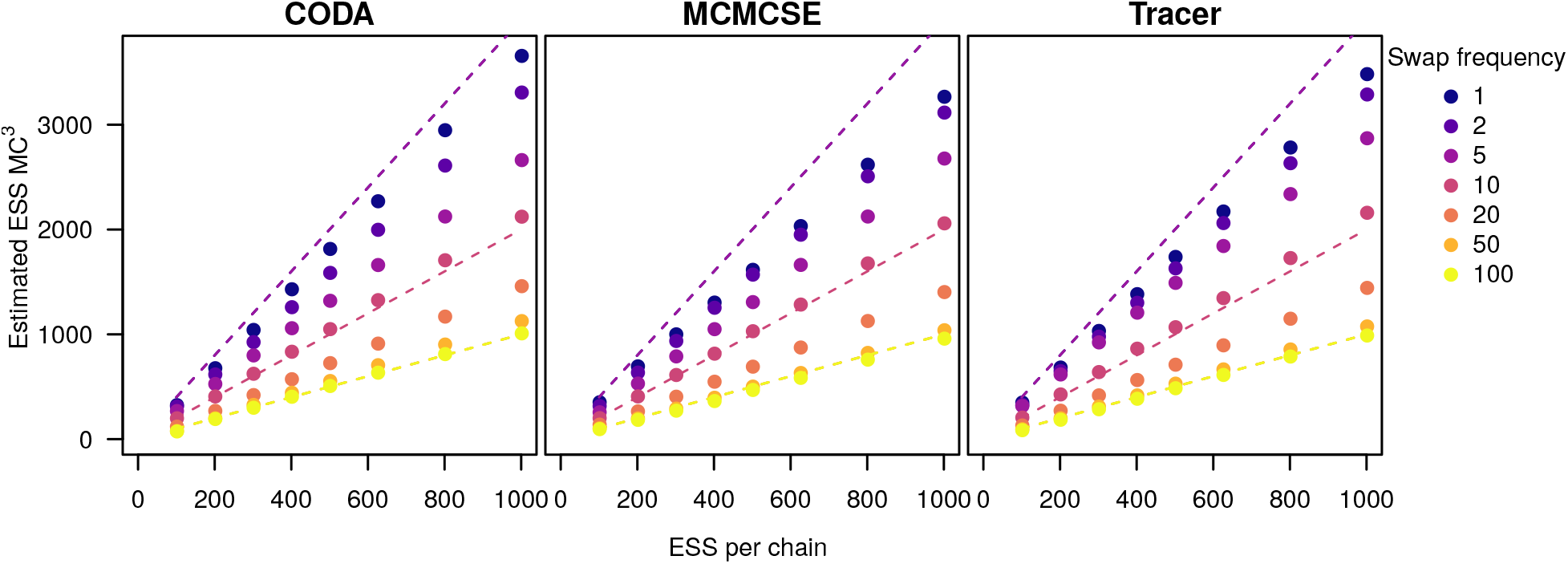
Evaluation of ESS estimation when samples are generated using our MC^3^ algorithm (Algorithm 3). The x-axis shows the true ESS used to generate the *K* = 4 different chains. The y-axis shows the estimated ESS value for samples from the MC^3^ chain. The colored dots show the average ESS estimate over 1,000 replicates. The dashed lines show the expectation for the ESS for each swap value, the expectations for the swaps of 1, 2 and 5 have equal values, thus they are represented by the same dashed line, the same happens for the swaps 20, 50 and 100.

##### Samples from adaptive MCMC

Almost all implementations of MCMC algorithms in phylogenetics and macroevolution use some type of adaptive MCMC (Ronquist *et al.*, 2012; Bouckaert *et al.*, 2014; Höhna *et al.*, 2016a,b). An adaptive MCMC algorithm changes the tuning parameter of the proposal distribution every κ iterations. For example, the window size of the sliding window proposal could be updated to achieve a target acceptance probability of 0.45 (Yang & Rodríguez, 2013; Höhna *et al.*, 2017). Thus, the autocorrelation time *τ* changes during the MCMC simulation. Tuning is performed during a pre-burnin phase where no samples are taken from the MCMC simulation or during the actual MCMC simulation if the tuning parameter is guaranteed to stabilize (i.e., converge) and tuning is performed with low frequency, e.g., *τ* < *κ*.

Here we only consider the second case where tuning is performed during the actual MCMC simulation while taking samples because the first case is equivalent to samples from a standard MCMC simulation. We sampled from 1,000 replicate MCMC simulations where the total chain length was broken into five intervals. During each interval, we generated samples from an MCMC simulation with *τ* = {50, 40, 30, 20, 10}, that is, in the first interval we had a higher autocorrelation and in the last interval we had the lowest autocorrelation. The size of the intervals varied between *κ* = {50, 100, 200, 500, 1000}. We expect a true ESS of

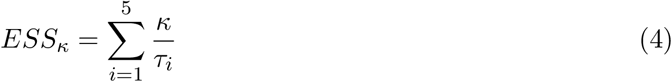

because in every interval *i* of size *κ* we should obtain 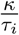 independent samples.

We observed that all three methods produce comparable estimated ESS values when samples are taken from an adaptive MCMC algorithm (Figure 3). However, for the adaptive MCMC samples `CODA` was slightly more conservative than `Tracer` and `MCMCSE` compared to our previous experiments. Furthermore, all three methods produced lower ESS estimates than our analytical expectation. This underestimation could be due to fact that all ESS methods compute an average autocorrelation time *τ* for all samples and not per-window estimates. Our observed estimated ESS values are closer to

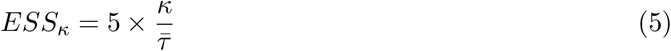

where 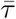 is the average autocorrelation time. Thus, ESS estimates are not inflated and instead conservative when samples are taken from adaptive MCMC algorithms.

**Figure 3:**
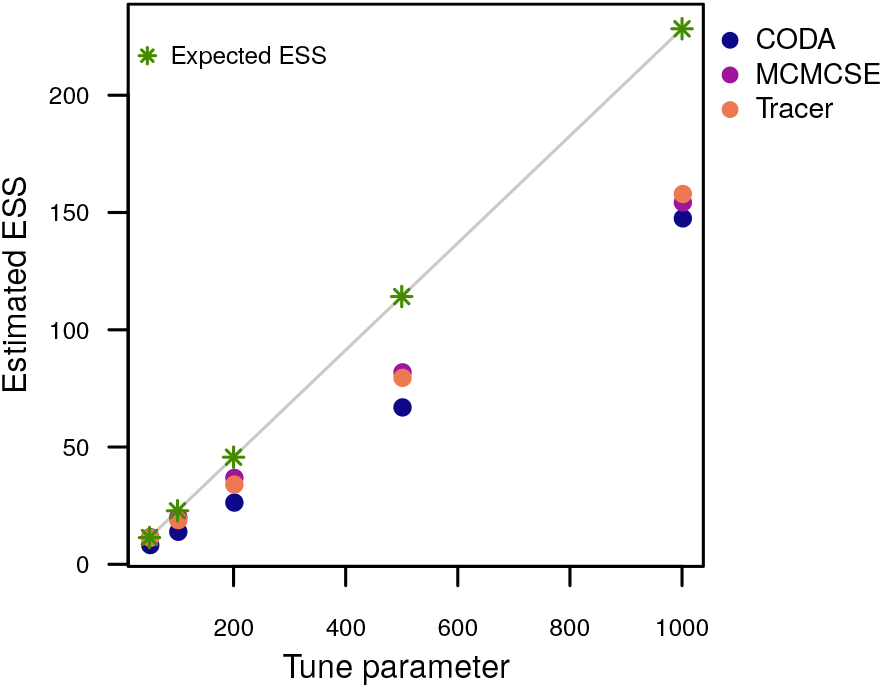
Evaluation of the estimated ESS for samples from adaptive MCMC simulations. The MCMC simulation was tuned every *κ* iterations in a total of four times. Within each of the five intervals we used an autocorrelation of *τ* = {50, 40, 30, 20, 10}. The x-axis shows the tuning frequency and the y-axis shows the estimated ESS. The colored dots show the average ESS of 1,000 replicates. The grey line shows the expected ESS values.

#### Assessing ESS estimation of discrete parameters

Tree topologies are arguably the most important but also most difficult parameter of Bayesian phylogenetic analyses (Harrington *et al.*, 2021). Tree topologies can be considered as categorical parameters and thus standard ESS estimation algorithms for continuous parameters do not apply. Therefore, we transform the samples of tree topologies into traces of absence/presence of splits. For each split that was sampled at least once during the MCMC simulation, we construct a trace that has a 1.0 if the split is present in the currently sampled tree and 0.0 otherwise. Hence, we obtain one discrete trace per split.

The three ESS estimation methods introduced above are derived for parameters drawn from continuous distributions. Whether these methods also work well for discrete, binary samples has not been studied. We approximate the sampled split frequencies using samples from a binomial distribution with probability *p* corresponding to the true posterior probability of the split. Different splits are clearly not independent because they are extracted from the same tree topologies; however, it suffices to consider only a single split when studying the behavior of the ESS estimation methods here.

Following our approach for continuous parameters, we simulated 1,000 replicates and sampled from our MCMC algorithm (Algorithm 1) with a true ESS of *N* = 625 (see Equation 2) and an autocorrelation of *τ* = {1, 5, 10, 20, 50}. We varied the true probability of the split *p* between 0.001 and 0.999 in increments of 0.001.

The efficiency of all methods dropped drastically once the true posterior probability gets close to either zero or one (Figure 4). This is expected because once the posterior probability is very close to zero or one, it is very likely that all samples either include or exclude the split. The ESS estimation methods are not designed to work well if all or all but one sample are zero (or one, respectively). It is intrinsically impossible to tell if all samples were identical because they are autocorrelated or because they truly should be the same. Thus, we suggest to exclude splits and parameters that have a posterior probability of *p* < 0.01 or *p* > 0.99.

**Figure 4:**
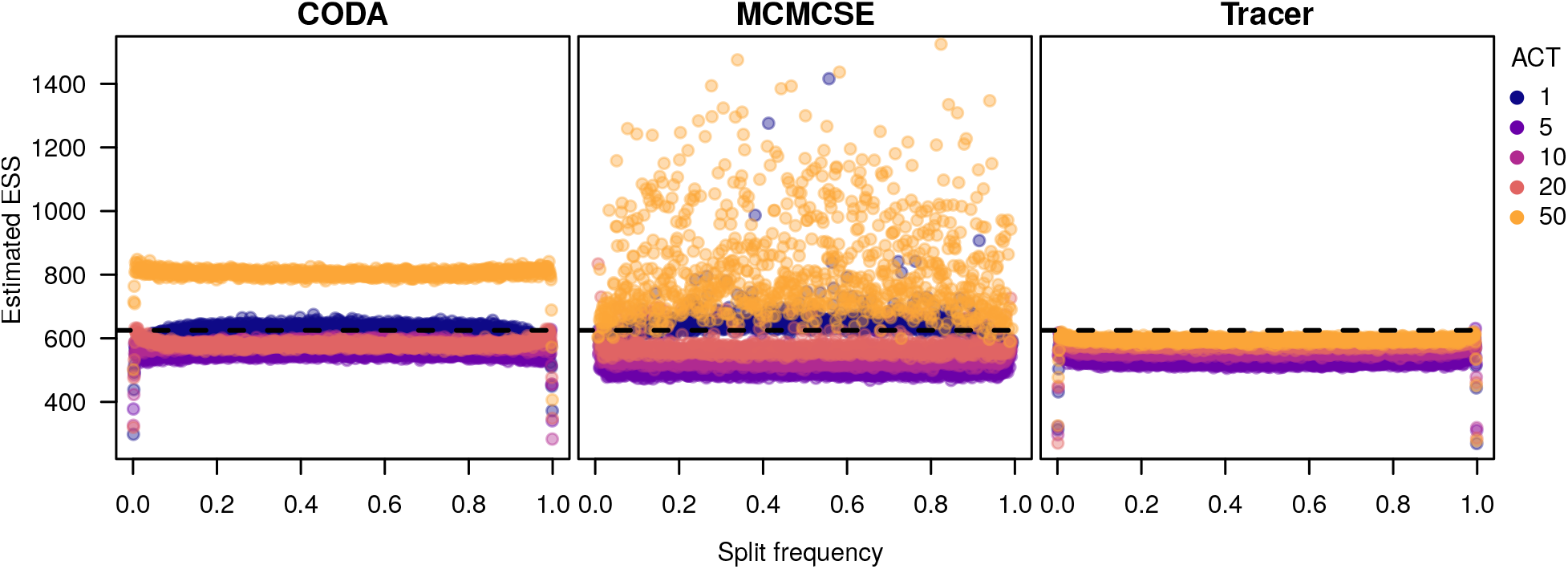
Evaluation of ESS estimation for samples taken from a binomial distributions with different autocorrelation times and varying probability *p*. The true ESS was *N* = 625 and *p* varied from 0 to 1 with steps of size 0.001. The colored dots show average ESS estimates for 100 replicates. The dashed line shows the true ESS value for the initial sample.

In conclusion, `Tracer` had an overall high precision in recovering the true ESS also when samples are binary (Figure 4). As before, we observed that `CODA` performs badly when the ACT was high. Similarly, `MCMCSE` performed poorly for an ACT of 50. Thus, `Tracer` is the only method we tested that is robust under all circumstances.

### REPRODUCIBILITY OF MCMC RUNS

In the previous section we focused on the precision of parameter estimates, which concerns the question of whether we have run the MCMC simulation long enough or if we need to run the MCMC simulation longer. In this section, we focus on the second question: “Are our parameter estimates *reproducible*?” Our estimates are reproducible if the MCMC simulation has converged to the true posterior distribution and did not get stuck in some other area of the parameter space. For example, single MCMC runs can achieve a high effective sample size but have sampled posterior probabilities that do no reflect the true posterior probabilities because the MCMC simulation got trapped in an “island in tree-space” (Höhna & Drummond, 2012; Whidden & Matsen, 2015).

In practice, we can never know with absolute certainty that our MCMC simulation has converged to the true posterior distribution. Instead, one commonly compares multiple independent MCMC runs. In phylogenetics, the two most commonly used approaches to compare independent MCMC runs are the potential scale reduction factor (PSRF, Gelman & Rubin, 1992) for continuous parameters and the average standard deviation of split frequencies (ASDSF, Nylander *et al.*, 2008; Lakner *et al.*, 2008) for tree topologies.

The PSRF computes the ratio of the variance of samples between chains over the variance of samples within a chain. If this ratio converges towards one, i.e., if the variance between independent chains is the same as the variance within a chain, then all MCMC simulation have presumably sampled from the same distribution and thus have converged to the true posterior distribution. However, the PSRF is problematic for two reasons: (1) there is no clear threshold and in practice values between 1.003 and 1.3 have been used (Vats & Knudson, 2020); and (2) the PSRF is very sensitive to the shape and variance of the posterior distribution and large independent samples (*N* > 1, 000) from the same distribution can yield a PSRF clearly larger than one (see Supplementary Figure S4). Thus, we discourage the use of the PSRF for assessing convergence of Bayesian phylogenetic MCMC simulations. Instead, we suggest to use the Kolmogorov-Smirnoff test (KS) to assess if two samples come from the same distribution (Brooks *et al.*, 2003). Below, we discuss and explore below the behavior of the KS test to assess convergence for continuous parameters.

The ASDSF computes the posterior probability of each sampled split in a Bayesian phylogenetic MCMC simulation. Then, the difference between the posterior probabilities per split for two runs are computed. We support the underlying idea of the ASDSF to break the sampled phylogenies into splits and use the frequencies of observing each split. However, computing the average difference between splits is problematic because (1) if the frequency of one split differs strongly (e.g., a frequency of one in the first run but zero in the second run) and all other splits have identical frequencies, then the ASDSF will be low enough to wrongly signal convergence; and the expected difference between two MCMC samples for the same split depends on the true posterior probability of the split (see below). We introduce our alternative version of the ASDSF below.

#### Assessing reproducibility of continuous parameter estimates

The two-sided KS test for two samples constructs the empirical cumulative distribution function of the samples. The test statistic *D* is the largest difference of the two empirical cumulative distribution functions *F*_1_ and *F*_2_, *D* = max_*x*_ |*F*_1_(*x*) − *F*_2_(*x*)|. The *D* statistic shows a significant departure from the expectation that both samples were drawn from the same distribution if 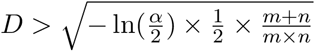 at a significance level *α*, assuming that the first sample has an ESS of *n* and the second sample an ESS of *m*.

If we would use the standard approach to define the threshold for *D* based on the number of samples, then we would reject on average *α* pairs of MCMC simulation regardless of how long we ran the MCMC simulations. To circumvent this problem, we fix *α* = 0.01 and *N* = 625 (see Equation 2) so that *D_crit_* = 0.0921. Thus, with more effective samples we should obtain a more precise estimate of *D* and therefore avoid incorrectly rejecting runs that had truly converged as often.

We assessed the false-rejection rate and power of the KS test with our threshold *D_crit_* by simulating 1,000 replicated pairs of samples with *N* = {100, 200, 625, 1000} *iid* values drawn from a normal distribution. The first normal distribution had a mean *μ*_1_ = 0 and standard deviation *σ* = 1. The mean of the second normal distribution differed by *μ*_2_ = {0.0, 0.04, 0.08, …, 0.8}, thus representing that the two means differed by 0% to 20% of the 95% probability interval.

When the samples where drawn from the same distribution (*μ*_1_ = *μ*_2_), then we observed a false rejection rate of 0.01 for a sample size of 625 (Figure 5). This rejection rate is exactly expected for an *α* = 0.01. If we increased the sample size to 1,000, then the false rejection rate decreased to 0.0003. Thus, more samples, i.e., longer MCMC runs, will increase the chances that the chains are assessed as converged if the samples are truly from the same distribution. When the mean of the distributions have a 10% difference (*μ*_2_ = *μ*_1_ + 0.1 × 4 × *σ*), then we correctly rejected convergence with a rate of 0.9974 for a sample size of 625 and 0.9995 for a sample size of 1,000.

**Figure 5:**
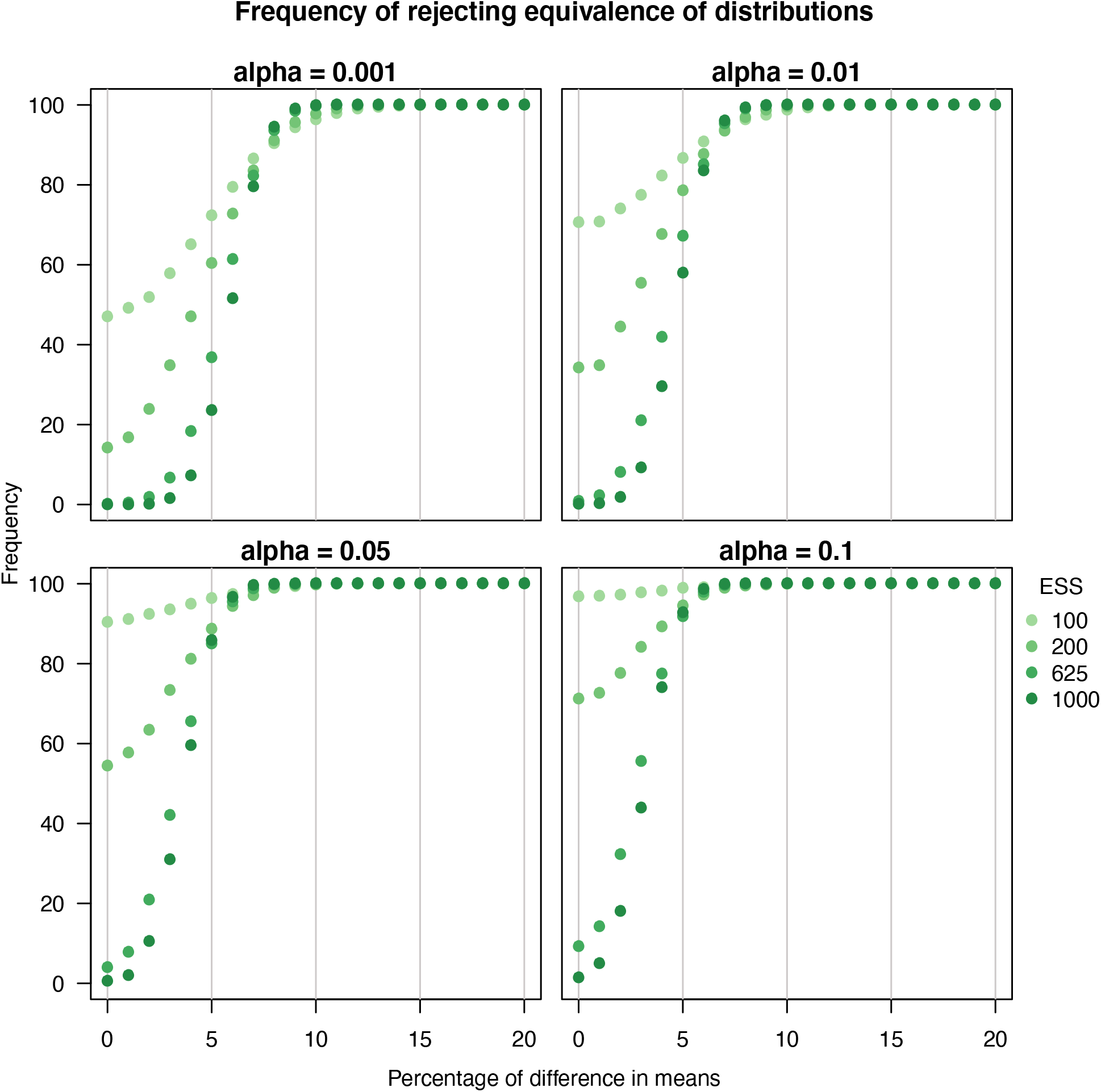
Testing the power of the Kolmogorov-Smirnov test statistic to distinguish between samples from two different distributions. The two samples were drawn from different normal distributions with different mean values. For each combination of difference in means, 1,000 replicates were tested and the frequency of rejecting the null hypothesis that both samples were drawn from the same distribution was computed (y-axis). The x-axis displays the difference in the means of the distributions with regard to the 95% probability interval of the normal distribution with mean 0 and standard deviation 1.

In the previous section we defined that our mean estimate is precise enough if the standard error is smaller than 1% of the 95% probability interval. Here we showed that the KS test has very strong power to reject runs if the true means of the samples was different by 10% or more, and has an acceptable power of 0.95 when the means are different by 8% (Figure 5). Increasing the ESS further would both decrease the standard error of the mean, i.e., increase our precision, and slightly increase the power to correctly reject convergence when the samples are truly from different posterior distributions.

The KS test is well established for testing if two samples are drawn from the same underlying distribution, which we also illustrated in Figure 5. However, the KS test is less established as a convergence assessment tool for autocorrelated samples drawn from MCMC simulations (but see Brooks *et al.*, 2003). Therefore, we explored if autocorrelation affects the behavior of the KS test. We performed the same test as before (1,000 replicates, two samples from normal distributions where the mean changed by *μ*_2_ = *μ*_1_ + *X* × 4 × *σ*) but introduced autocorrelation of *τ* = {1, 5, 10, 20, 50} into the samples using Algorithm 1. We observed that there is no impact of using autocorrelated or uncorrelated samples for the false rejection rate and power of the KS test (Figure 6 for *N* = 625).

**Figure 6:**
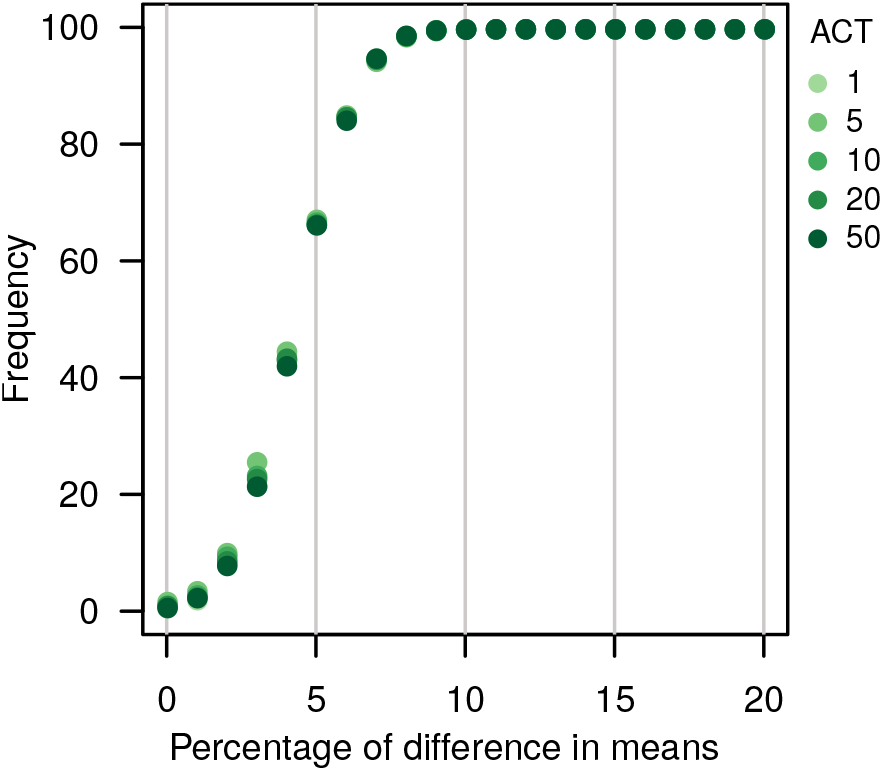
Kolmogorov-Smirnov test to detect difference in distribution for autocorrelated samples. The two samples were drawn from different normal distributions with different mean values and different ACT values. Here we only show the results for *N* = 625. For each combination of difference in means, 1,000 replicates were tested and the frequency of rejecting the null hypothesis that both samples were drawn from the same distribution was computed (y-axis). The x-axis displays the difference in the means of the distributions with regard to the 95% probability interval of the normal distribution with mean 0 and standard deviation 1.

#### Assessing reproducibility of discrete parameter estimates (split frequencies)

Phylogenetic trees and split frequencies do not yield continuous distributions which can be compared using the KS test. Instead, one often plots the split frequencies of one run against a second run (xy-plot). If there is a strong deviation from the diagonal line, then the two runs are assessed as non-converged. Quantitatively, one can compute the average (Lakner *et al.*, 2008) or maximum (Höhna & Drummond, 2012) deviation deviation of split frequencies. However, an under-appreciated characteristic is that the expected difference, or standard error, of split frequencies depends strongly on the true split frequency.

In our view, the ASDSF is not sensitive enough to detect outliers in estimated split frequencies. For large trees with many splits, there will be many splits with a very low frequency. These splits overwhelm the computation of ASDSF and outliers, such that even a difference in posterior probability as large as 1.0 in one run and 0.0 in the second might not be detected. Another unsolved issue with the ASDSF is that no theory for a threshold is provided and the default thresholds are applied to all tree sizes. Additionally, the difference for each split has equal weight although the stochastic difference in split frequencies depends on the true split frequency.

Samples of split frequencies can be treated as a series of 1.0 and 0.0 (presence and absence, as we did above). Thus, we can consider samples of split frequencies as draws from a binomial distribution. For a binomial distribution, we can actually calculate the expected difference between two samples analytically. Let us denote the expected difference between two samples for a split with true frequency *p* as 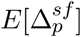. Then, if each sample has the size *N*, we can simply sum over all possible outcomes weighted by their difference in splits, which is

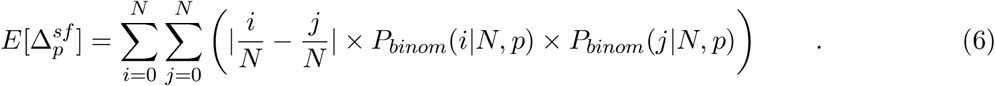

Figure 7 shows curves of the expected difference in split frequencies 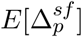, EDSF, for different samples sizes. As expected, 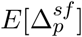 is smallest when the true split frequency is close to the boundaries zero or one, and largest when the true split frequency *p* = 0.5. Moreover, 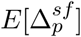 decreases for larger sample sizes. Interestingly, for all possible true split frequencies we expect at most a difference of 0.014 (or 0.026 and 0.04) if we have 625 independent samples (or 200 and 100 respectively). For our purposes, we define that two samples of phylogenetic trees are from the same underlying posterior distribution if all splits have difference smaller than the 95% quantile of the 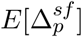 for *N* = 625 and 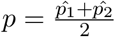, where 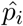 is the estimated split frequency for run *i*. By using the 95% quantile of the 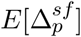, we guarantee that only 5% of the difference in splits will be rejected when they are truly coming from the same posterior distribution.

**Figure 7:**
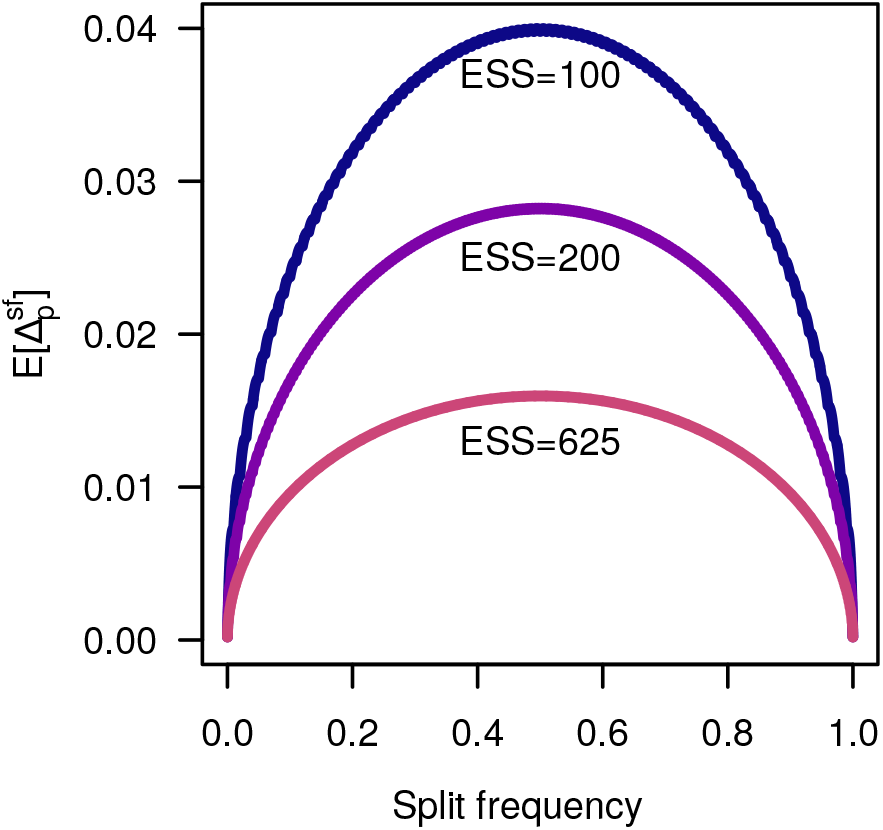
The expected difference in split frequencies for ESS of 100, 200 and 625. The x-axis is the true value of the split frequency. The y-axis is the expected difference in split frequencies. The effect of increasing the ESS is the decrease of differences in frequency of sampled splits.

### Convenience: Implementation of convergence assessment and interpretation of output

We implemented the methods described here in the stand-alone R package Convenience. `Convenience` is open-source and can be downloaded and installed from https://github.com/lfabreti/convenience. Currently, `Convenience` supports the output file formats from `RevBayes` (Höhna *et al.*, 2016b), `MrBayes` (Ronquist *et al.*, 2012), `BEAST` (Bouckaert *et al.*, 2014) and `PhyloBayes` (Lartillot *et al.*, 2009). Additionally, it is also possible to assess convergence in outputs containing only continuous or discrete parameters (trees).

The main function of Convenience is `checkConvergence`(), which runs the complete convergence assessment pipeline and the thresholds established in this article.

~~~
> test_convergence <− checkConvergence (“convenience /example/”)
~~~

The `checkConvergence()` function checks first the best burn-in value. If the burn-in is greater than 50%, the function stops and tell the user that the burn-in is too large. Otherwise, the function continues the convergence assessment by applying the described methods to the continuous and discrete parameters. In addition, the user has the possibility to use each method separately in different functions and change the thresholds as suited. A more detailed explanation of the functions can be found in the tutorial at https://revbayes.github.io/tutorials/convergence/.

Once the convergence assessment has been performed, the user has the option to print or plot the results. `Convenience` produces four main plots:

~~~
> plot Ess Continuous (test_convergence)
> plot Ess Splits (test_convergence)
> plotKS (test_convergence)
> plot DiffSplits (test_convergence)
~~~

`plotEssContinuous` displays the ESS values for all continuous parameters and all MCMC replicates within one plot (Figure 8a). If MCMC convergence has been achieved, then all ESS values in this histogram are on the right side of the minimum ESS threshold. `plotEssSplits` displays the ESS values for all splits and all MCMC replicates (Figure 8b). Again, if MCMC convergence has been achieved, then all ESS values in this histogram are on the right side of the minimum ESS threshold. `plotKS` displays the KS-score for all continuous parameters and all pairwise comparisons of MCMC replicates (Figure 8c). If MCMC convergence has been achieved, then all KS scores in this histogram are on the left side of the maximum KS threshold. `plotDiffSplits` displays the difference in split frequencies for all splits and all pairwise comparisons MCMC replicates (Figure 8d). If MCMC convergence has been achieved, then all differences in split frequencies are below the maximum split frequency threshold.

**Figure 8:**
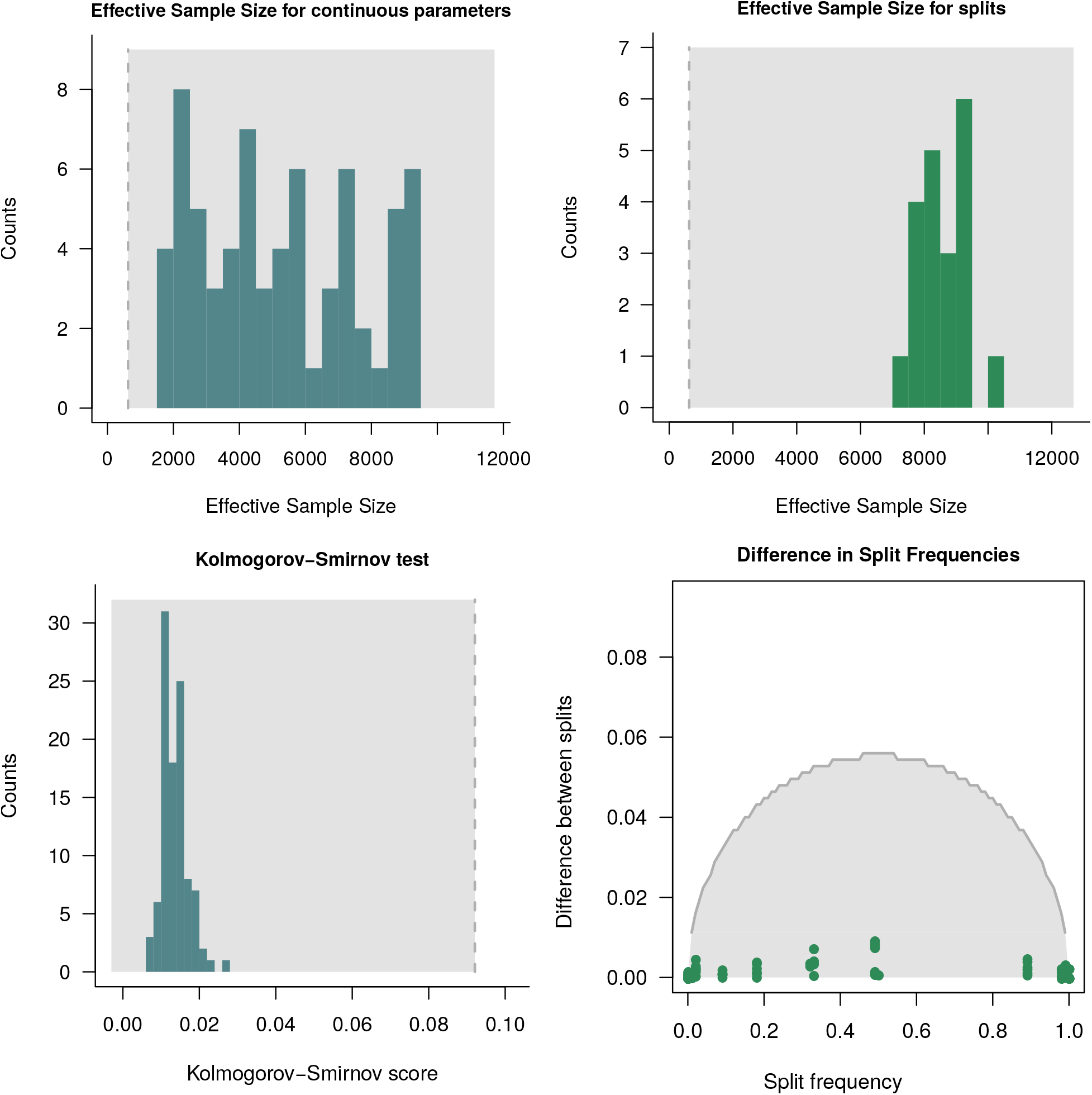
The plots generated with `Convenience` for summarizing and visualizing the results from the convergence assessment. Here we used the MCMC example in the `RevBayes` tutorial https://revbayes.github.io/tutorials/ctmc/, see Höhna *et al.* (2017). Top-left: the histogram of estimated ESS values for the model parameters (continuous parameters). Top-right: the histogram of calculated ESS for the splits. In both histograms the dashed lines represents the minimum ESS threshold of 625. Bottom-left: histogram of the Kolmogorov-Smirnov (KS) test for the model parameters, the dashed line represents the threshold for the KS test. Bottom-right: the observed difference is split frequencies in the green dots and the maximum threshold for split frequencies based on the expected difference between split frequencies (EDSF) in the gray curve. For all plots the gray area shows where the values should be if the analysis achieved convergence.

In summary, applying `Convenience` is fully automatic. Thus, the package can be used interactively or in batch-mode (e.g., on computer clusters). If an MCMC analysis includes a plot similar to Figure 8, then it is easy to verify convergence assessment. The text output of Convenience can be easily parsed to perform hundreds or thousands of convergence assessments.

## Discussion and conclusions

### CONVERGENCE THRESHOLDS FOR NUISANCE PARAMETERS

In many phylogenetic analyses, there are some focal parameters and other parameters are nuisance parameters. For example, in a traditional phylogeny estimation analysis, the phylogenetic tree is the focal parameter and the substitution model parameters might be nuisance parameters. So far, the thresholds used in convergence assessment are applied equally to all parameters. That is, one requires that all parameters have an ESS> 625 (or whichever other threshold was used). One might argue that checking for convergence for the nuisance parameters is not as relevant as checking for convergence for the focal parameters. Thus, it could be possible to use more relaxed thresholds for the nuisance parameters. However, we find no theoretical support for treating nuisance and focal parameters differently. Whether relaxing the precision, and consequently the ESS, for the nuisance parameters can affect the convergence of the focal parameters needs to be further investigated. For now, we advise on using the same criteria for all underlying parameters of the model.

### FUTURE DIRECTIONS

Our approach and evaluation presented here has several limitations and is only another small step towards robust and automatic convergence assessment. First, we did not test how well either of these methods perform when the posterior distribution is multi-modal. Second, parameter non-identifiability might confuse convergence assessment tests. For example, hidden Markov models can produce seemingly multi-modal posterior distributions if the hidden state is arbitrarily labelled and not ordered (Lartillot *et al.*, 2007; Baele *et al.*, 2021). Third, we currently reject convergence if a single MCMC run did not converge or produced a different posterior distribution. When MCMC mixing is very challenging, it might happen that many replicated MCMC runs are performed and only a small subset converged. Thus, it would be desirable to automatically identify the subset of MCMC runs that converged.

### CONCLUSIONS

Convergence assessment should be a mandatory, objective, simple and reproducible step in any Bayesian analysis that relies on samples from the posterior distribution. In this manuscript we presented and explored our approach, which is implemented in the R package `Convenience`. We identified two crucial aspects when running an MCMC simulation: (i) Has the MCMC ran long enough?; and (ii) Do the samples represent the true posterior distribution?

We addressed the first question by focusing on the precision of the posterior mean estimate. If we have sufficiently many samples from the posterior distribution, then our standard error of the mean estimate will be sufficiently small. Thus, one only needs to check if the effective sample size is large enough and we provide some objective criteria to choose a threshold for the minimum ESS. If we accept an SEM of 1% of the 95% credible interval, then a minim ESS of 625 is required. We tested three commonly used methods to estimate the ESS: spectral density estimators of an auto-regressive process (`CODA`), batch means (`MCMCSE`), and initial sequence estimator (`Tracer`). Our assessment included: (a) independent samples; (b) autocorrelated samples; (c) samples from Metropolis-Coupled MCMC simulations; and (d) samples from adaptive MCMC simulations. We found that only the initial sequence estimator (`Tracer`) was robust in all scenarios and for all ranges of autocorrelations.

Focusing on phylogenetic applications, we showed that samples from the posterior distribution of phylogenetic trees can be converted into binary traces of absence/presence of splits. The ESS estimation works robustly on these discrete, binary traces and can be applied in the same way.

We addressed the second question by focusing on reproducibility of multiple MCMC runs. We observed that the commonly used potential scale reduction factor (PSRF) is not robust to the shape of the posterior distribution. For example, samples from a lognormal distribution yield a PSRF that is asymptotically significantly larger than 1.0. We suggest the Kolmogorov-Smirnov test instead, which we showed to work well also for autocorrelated samples.

We modified the average standard deviation of split frequencies (ASDSF) to use instead an analytically derived expected difference between split frequencies (EDSF). We demonstrated that the EDSF depends on the true frequency of a split, and thus the same thresholds for all splits cannot be used.

