## Supplementary Material for "Convergence Assessment for Bayesian Phylogenetic Analysis using MCMC simulation"

### Convergence Assessment in Phylogenetics

### Contents

**S1 Precision of an estimator to assess sufficiently many samples** **3**

**S2 ESS Estimates for Independent Monte Carlo Samples** **4**

**S3 ESS Estimation of Samples from a Binomial Distribution** **5**

**S4 Potential Scale Reduction Factor (PSRF)** **6**

**S5 Estimating burn-in length** **8**

#### S1 Precision of an estimator to assess sufficiently many samples

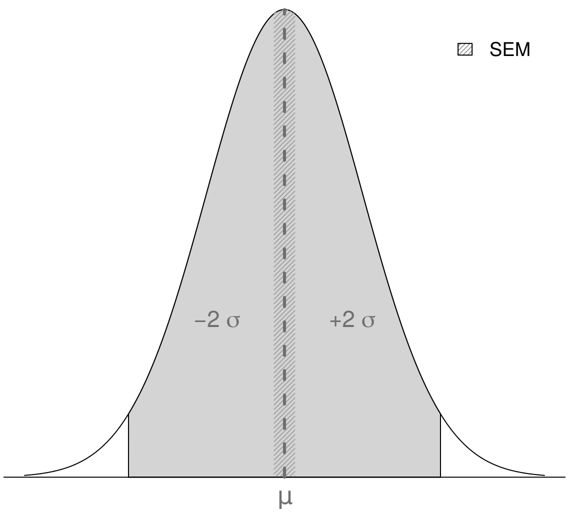


**Figure S1:** Schematic of a posterior distribution with mean estimate $\mu$ and 95% credible interval. Here, we chose a normal distribution to represent the posterior distribution. The shaded area shows the 95% credible interval, which has a width of $\pm$ 2 $\sigma$. The dashed line represents the mean ($\mu$) of the distribution. The darker shaded area represents the standard error of the mean. If the SEM is smaller or to 1% of the 95% credible interval size, then we accept the mean estimate as sufficiently precise.

#### S2 ESS Estimates for Independent Monte Carlo Samples

We assessed the ability of `CODA`, `MCMCSE` and `Tracer` to estimate the true ESS from an independent sample. This represents the case where we thinned our MCMC samples and all values are virtually independent. Furthermore, in this first test case we were interested in whether the shape of the distribution had an impact on the efficiency of the method. Thus, we used a normal distribution with mean 0 and standard deviation 1, a lognormal distribution with mean 0 and standard deviation 1 (both on the log scale) and an exponential distribution with rate 1. For each distribution we simulated 1,000 replicates of $N = \{100, 200, 300, 400, 500, 625, 800, 1000\}$ independent and identically distributed ($iid$) samples (i.e., $N$ is equal to the true ESS). Then, we estimated the ESS of these simulated samples using `CODA`, `MCMCSE` and `Tracer`.

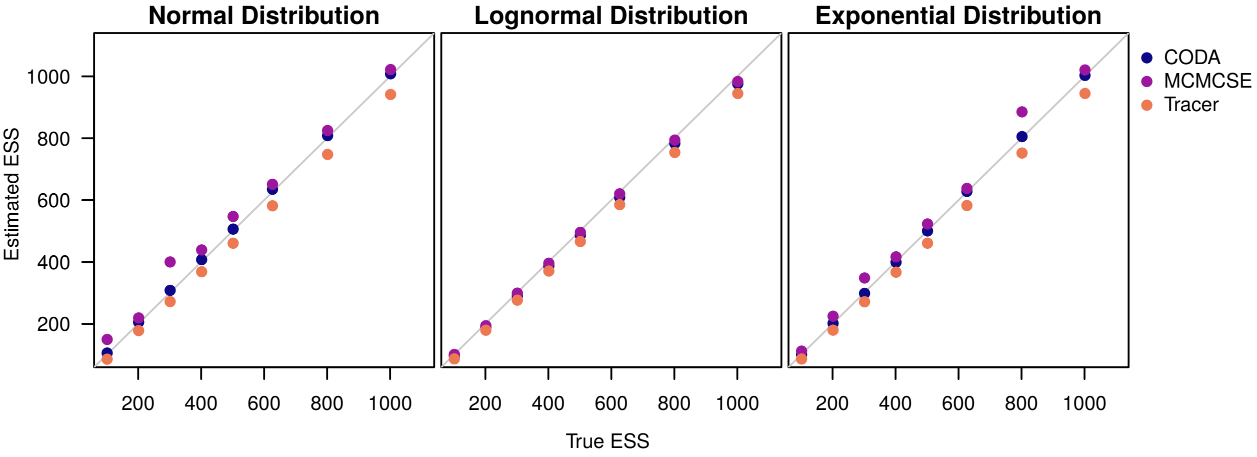


**Figure S2:** Estimated Effective Sample Sizes (ESS) for independent samples from different continuous distributions. We used `CODA`, `MCMCSE` and `Tracer` to estimate the ESS and evaluate their accuracy. The x-axis is the true ESS used to generate the sample, the y-axis is the average estimated ESS. The samples were simulated under a normal, lognormal and exponential distribution (from left to right).

All three methods perform comparably well and appear sufficiently precise and robust (Figure S2). Neither the number of samples nor the choice of the shape of the underlying distribution impacted the accuracy. Thus, we will only use the normal distribution in the following experiments. `Tracer` always had the lowest estimated ESS and thus is the most conservative of the three methods. `CODA`, on the other hand, had the highest overall precision.

#### S3 ESS Estimation of Samples from a Binomial Distribution

Figure S2 shows the estimation of the ESS for a normal, lognormal and exponential distribution. The same tests were performed for the binomial distribution for different $p$ and the results are shown in Figure S3. For each value of $p$ and sample size, we replicated 1000 test to calculate the mean estimated ESS. The estimation of the ESS is not influenced by the probability of success($p$) from the binomial distribution, since all plots in Figure S3 exhibit the same tendency.

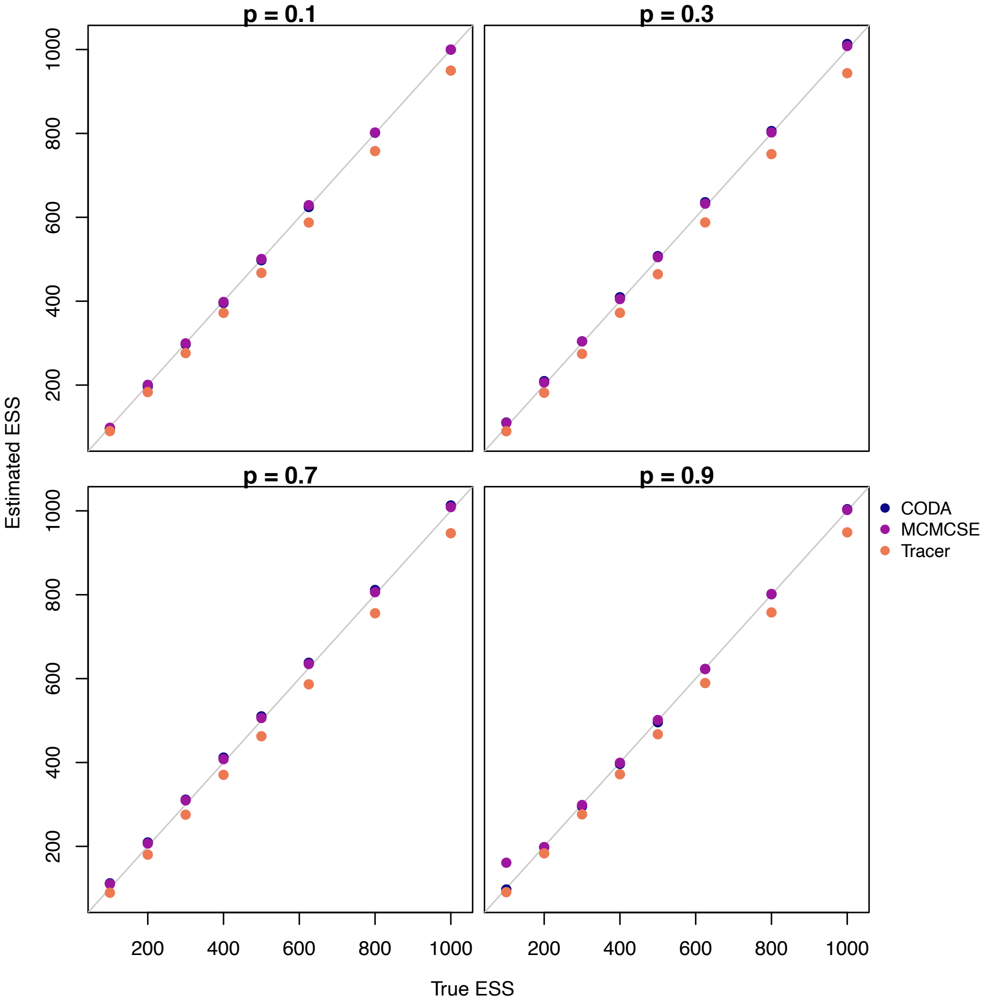


**Figure S3:** The estimated Effective Sample Size (ESS) for samples drawn from a binomial distribution with different probabilities of success ($p$). The $p$ is indicated on top of each plot. For each plot we varied the size of the sample drawn (true ESS) from 100 to 10000. We then calculated the ESS with three methods: CODA, MCMCSE and Tracer. The blue dots correspond to CODA, the purple dots to MCMCSE and the orange dots to Tracer.

### S4 Potential Scale Reduction Factor (PSRF)

In phylogenetics, the only (or at least most widely) used convergence assessment method for continuous parameters between replicated MCMC runs is the potential scale reduction factor [PSRF, 1]. We tested the PSRF under different common statistical distributions (normal distribution, gamma distribution and lognormal distribution). In practice, we never know the true shape of the posterior distribution when using empirical data. It is possible that the posterior distribution is well approximated by a normal distribution, but it is similarly possible that the posterior distribution is skewed and thus better approximated by a lognormal or gamma distribution.

We generated two sets of values, according to the same distribution, to compare the performance of evaluating the variance within each set and between sets. Each set had $1 \times 10^5$ values drawn from the specified distribution. The distributions used were the normal, lognormal, exponential and gamma. For each distribution, we used different values of variance (1.0, 4.0, 25.0, 100.0) to compare the behavior of the PSRF when the variance of the set of values changes. For the normal and exponential distributions the PSRF values varied from 1.001 to 1.019 as we increased the size of the samples. The values for both distribution had a maximum difference of 0.01 and increasing the variance of the samples did not affect the estimated PSRF values. In the tests for the lognormal and gamma distribution, the PSRF values varied according to the variance of the sample, as it is shown in Figure S4

Surprisingly, we observed that the PSRF does not converge towards 1.0 with increasing sample size if the distribution is heavily skewed (Figure S4). For example, when we simulated values from a gamma distribution with $\theta = 10$ and varied $\kappa$, PSRF values were comparably high (PSRF $> 1.05$). Similarly, when we used simulated values from a lognormal distribution, PSRF values never converged towards 1.0 and the asymptotic PSRF increased with higher variances. Thus, we conclude that the PSRF is not universally applicable in phylogenetics and other approaches, such as the Kolmogorov-Smirnov test, are superior.

Figure S4 shows the PSRF values calculated for 3 different distributions: normal, lognormal and exponential. The mean PSRF was calculated by comparing $1 \times 10^5$ times two sets of values drawn from the same distribution with equal parameters. We used 4 different variance values (1.0, 2.0, 5.0 and 10.0) for each distribution to compare how they affect the PSRF estimate.

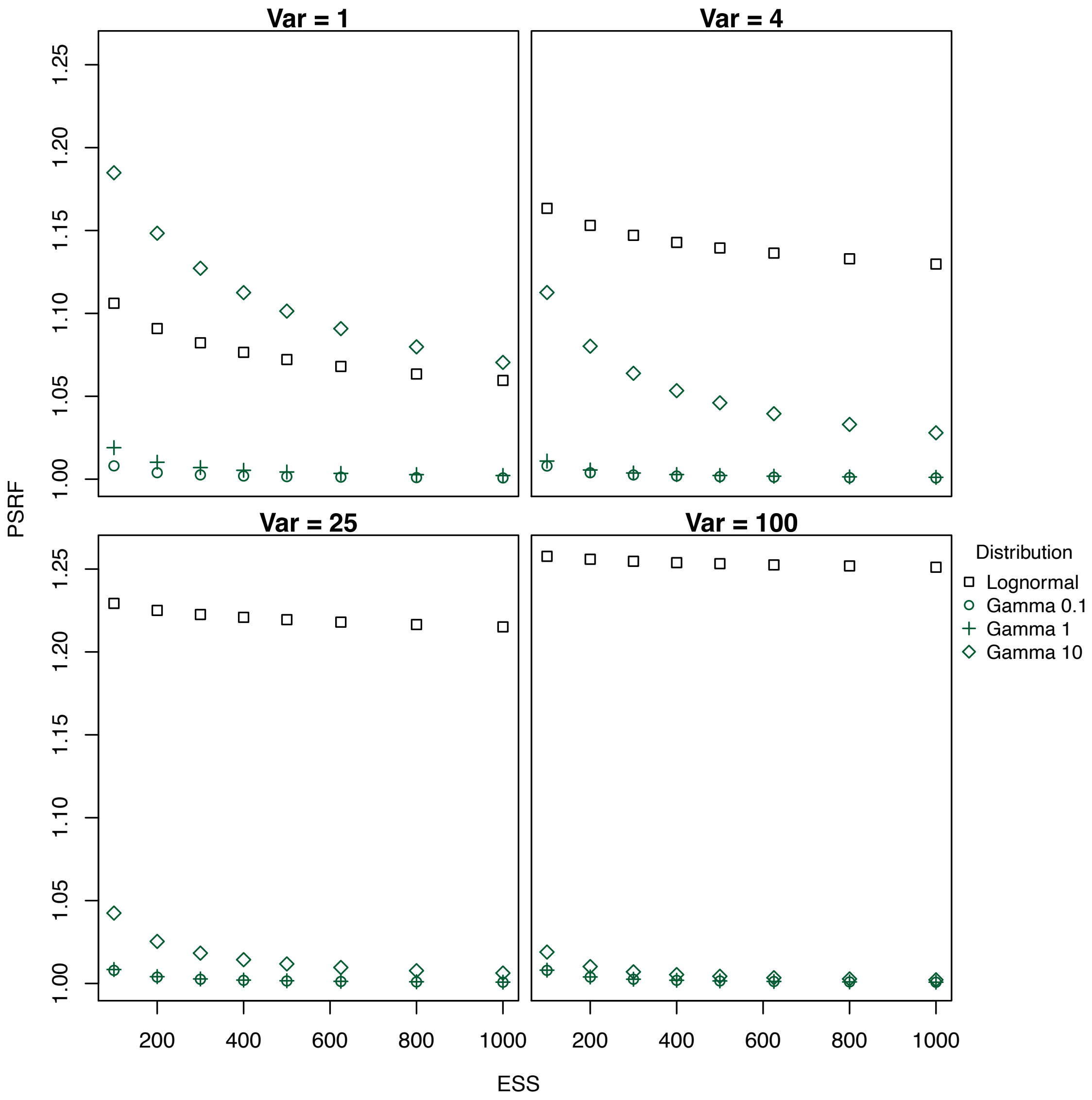


**Figure S4:** The potential scale reduction factor (PSRF) values calculated for two samples drawn under the same distribution. The variance of the distributions assumed values of 1, 4, 25 and 100, as is shown on top of each plot. The plotted values correspond to the mean of $1 \times 10^5$ replicates. The black squares represent the values calculated from a lognormal distribution. The green figures correspond to the Gamma distribution. The circle, the cross and the diamond correspond to scales parameters of 0.1, 1 and 10, respectively.

### S5 Estimating burn-in length

Most recorded samples from an MCMC algorithm in phylogenetics do not start with a random draw from the posterior distribution. Instead, most MCMC algorithms are either initialized with fixed starting values [2, 3] or random values drawn from the prior distribution [4]. It is therefore necessary to remove the first $X\%$ of samples as burn-in to obtain an unbiased approximation from the posterior distribution. In phylogenetics, a common burn-in is either 10% or 25%, depending on the arbitrary preference of the software developer.

To make the burn-in selection slightly less arbitrary, we developed the following procedure. We search for the optimum burn-in value defined as the lowest burn-in that passes the convergence tests. We start with no burn-in and increase it by 10% up to a maximum of 50% of the chain. If more than 50% of the chain has to be discarded, the MCMC spent too much time outside of the stationary distribution and the whole analysis should be redone and/or run longer.

**Algorithm S1** Estimating the burn-in length.

```
Inputs:
    X: the samples.

Initialize:
        n ← length(X)                          ▷ the total number of correlated samples

for i in {0.0, 0.1, 0.2, 0.3, 0.4, 0.5} do                 ▷ generate independent chains
    Y ← X[(i∗n):n]                                ▷ retrieve the post-burnin samples
    if convergence(Y) == pass then                      ▷ check for convergence
        break                                                            ▷ stop
    end if
end for

return i
```
